# Circular mitochondrial-encoded mRNAs are a distinct subpopulation of mitochondrial mRNA in *Trypanosoma brucei*

**DOI:** 10.1101/2023.02.10.528059

**Authors:** Clara M. Smoniewski, Poorya Mirzavand Borujeni, Austin Petersen, Marshall Hampton, Reza Salavati, Sara L. Zimmer

**Affiliations:** Department of Biomedical Sciences, University of Minnesota Medical School Duluth Campus, Duluth, MN, USA; Institute of Parasitology, McGill University, Quebec, Canada; Department of Biology, University of Minnesota Duluth, Duluth, MN, USA; Department of Mathematics and Statistics, University of Minnesota Duluth, Duluth, MN, USA

**Keywords:** kinetoplastid, neglected tropical disease, trypanosomiasis, RNA degradation, polyadenylation, uridylation, RNA ligase

## Abstract

Since the first identification of circular RNA (circRNA) in viral-like systems, reports of circRNAs and their functions in various organisms, cell types, and organelles have greatly expanded. Here, we report the first evidence of circular mRNA in the mitochondrion of the eukaryotic parasite, *Trypanosoma brucei*. While using a circular RT-PCR technique developed to sequence mRNA tails of mitochondrial transcripts, we found that some mRNAs are circularized without an *in vitro* circularization step normally required to produce PCR products. Starting from total *in vitro* circularized RNA and *in vivo* circRNA, we high-throughput sequenced three transcripts from the 3’ end of the coding region, through the 3’ tail, to the 5’ start of the coding region. We found that fewer reads in the circRNA libraries contained tails than in the total RNA libraries. When tails were present on circRNAs, they were shorter and less adenine-rich than the total population of RNA tails of the same transcript. Additionally, using hidden Markov modelling we determined that enzymatic activity during tail addition is different for circRNAs than for total RNA. Lastly, circRNA UTRs tended to be shorter and more variable than those of the same transcript sequenced from total RNA. We propose a revised model of Trypanosome mitochondrial tail addition, in which a fraction of mRNAs is circularized prior to the addition of adenine-rich tails and may act as a new regulatory molecule or in a degradation pathway.

## Introduction

The presence of organelles is a distinguishing feature of all eukaryotes. Of the organelles evolutionarily acquired by symbiosis and thus retaining independent genomes, the mitochondrion exists in nearly all extant eukaryotes. The pared-down mitochondrial genome generally retains a remarkably similar core subset of a few coding loci and rRNAs across eukaryotic species. However, the overall organization of these mitochondrial genomes, and the mechanisms for expression and processing of their RNA products vary widely.

Circular RNAs (circRNAs) are a class of RNAs in which the 5’ and 3’ ends of a single RNA molecule are covalently ligated. They were first reported as viroids [1] but soon after discovered in the cytoplasm of HeLa cells [2] and yeast mitochondria [3]. Once thought rare, circRNAs’ ubiquity in nature is being increasingly recognized [4–6]. The precise mechanism of ligation remains an active area of research, and different mechanisms may operate in different cellular contexts. Ribozymes, cellular ligases, and spliceosomes have all been shown to mediate RNA circularization [7]. Most identified circRNAs are formed from eukaryotic mRNA introns or exons spliced at normal splice sites and subsequently ligated [6]. However, a 2018 report revealed the presence of circRNAs formed from full-length mitochondrial mRNAs in the unicellular alga *Chlamydomonas reinhardtii* [8]. The mitochondrial mRNA of this alga, like that of a mammalian cell, lacks a 5’ UTR and thus a ribosome binding site. The authors speculated that circularization of *Chlamydomonas* mitochondrial mRNAs provided a mechanism for ribosome binding upstream of the translational start site. The same group subsequently explored whether such a mechanism might also exist in the mammalian mitochondrion [9]. This was determined to be unlikely, because while circRNAs derived from *Chlamydomonas* mitochondrial mRNAs were full length with a termination codon, those from mammalian cells were truncated and thus non-translatable. Very recently, widespread mitochondrial circRNAs originating from mRNAs were discovered in plants, and some were shown to be ribosome associated [10]. The possibility that diverse regulatory roles for mitochondrial circRNAs exist has recently been confirmed with studies demonstrating that mitochondrial circRNAs regulate protein import into the mitochondria of metazoans and the output of mitochondrial reactive oxygen species in human cells [11,12]. Despite these few discoveries of functional roles for mitochondrial circRNAs, their functions and their associated mechanisms remain largely elusive. Further, the landscape of mitochondrial species of circRNAs among eukaryotic species across the tree of life has been barely scratched. The possibility of functional mitochondrial circRNAs in less-studied eukaryotic species is intriguing.

*Trypanosoma brucei* is an insect-transmitted parasitic protozoan of mammals that is part of a larger group called the kinetoplastids. These flagellates possess one unique mitochondrion per cell with a concatenated network of DNA and associated proteins called the kinetoplast. The kinetoplast (k)DNA contains 20 genes: 2 rRNAs and 18 mRNAs that mostly code for proteins of mitochondrial electron transport chain complexes. The process of kDNA transcription into RNA and subsequent RNA maturation steps have been extensively studied and a working model has emerged [13]. Evidence suggests that transcription is monocistronic [14], contrary to previously held hypotheses of polycistronic transcription [15], but that there may be a degree of both cannot be ruled out. After transcription, *T. brucei* RNAs may possess a 5’ triphosphate, which on mRNAs is converted to a 5’ monophosphate by the 5’ pyrophosphohydrolase complex (PPsome) [14]. All RNA is processed by the mitochondrial 3’ processome (MPsome), which trims the 3’ end and can add oligouridine (U) tails [16]. mRNAs generally have short adenine (A)-rich A/U heterogenous tails added by kinetoplast poly(A) polymerase I (KPAP1) as part of the kinetoplast polyadenylation complex (KPAC) in combination with KRET1 of the MPsome. Interactions between the 5’ and 3’ bound protein complexes are hypothesized to create a molecule with protected 5’ and 3’ ends in close proximity and are a key regulatory step against mis-tailing and extensive 3’ trimming [13]. These interactions are thought to be mediated and possibly coordinated by the RNA editing substrate-binding complex (RESC) [13]. The RESC is one of the main protein complexes that is involved in a complicated editing process that takes place for 12 of the 18 mRNAs. This unique uridine insertion/deletion RNA editing requires a multitude of proteins, many of which are found in the RESC and the RNA editing catalytic complex (RECC). After editing is complete, the short A-rich mRNA tail is elongated to a more heterogenous A/U tail before translation takes place. A multitude of RNA binding proteins control this process, including 3’ trimming, tailing, and editing.

Here, we present evidence of mitochondrial mRNA-derived circRNAs in the kinetoplastids that are very evolutionarily distant from both plants and metazoans. Mitochondrial circRNA of multiple mRNA transcripts are present in both the insect and mammalian bloodstream life stage forms of *T. brucei*. Using circular RT-PCR and deep sequencing, we confirmed that the 3’ and 5’ ends of these mRNAs are covalently linked together. We have investigated trends in the UTRs and tails that these transcript populations contain and show that for each transcript examined, its circRNAs are a distinct subpopulation of total mRNA that possess different tail and 5’ UTR characteristics. We hypothesize that the differences between a transcript’s linear and circular mRNA populations are an indication of an alternative pathway in the mRNA lifecycle differing from its normal maturation and turnover pathways.

## Results

### Some mitochondrial mRNA are circularized

CircTAIL-seq is a technique used to Illumina sequence individual transcript PCR libraries of mitochondrial mRNA tails. The approach captures enough of each molecule’s 5’ and 3’ termini that the presence of editing can be confirmed if necessary [17]. Library preparation first requires circularizing total RNA with T4 RNA ligase prior to the generation of cDNA, after which outward facing primers (“Circular primers” in **Figure 1A)** can generate PCR products capturing ligated 5’-3’ junction sequence (termed ligation point junction sequences). Individual sequence reads thus consist of the 3’ end of a transcript, its 3’ UTR, any 3’ tail, 5’UTR, and the start of its coding region. Unlike traditional inward facing primers, circular primers will not produce a product unless the transcript is first circularized (**Figure 1A)**.

**Figure 1.**
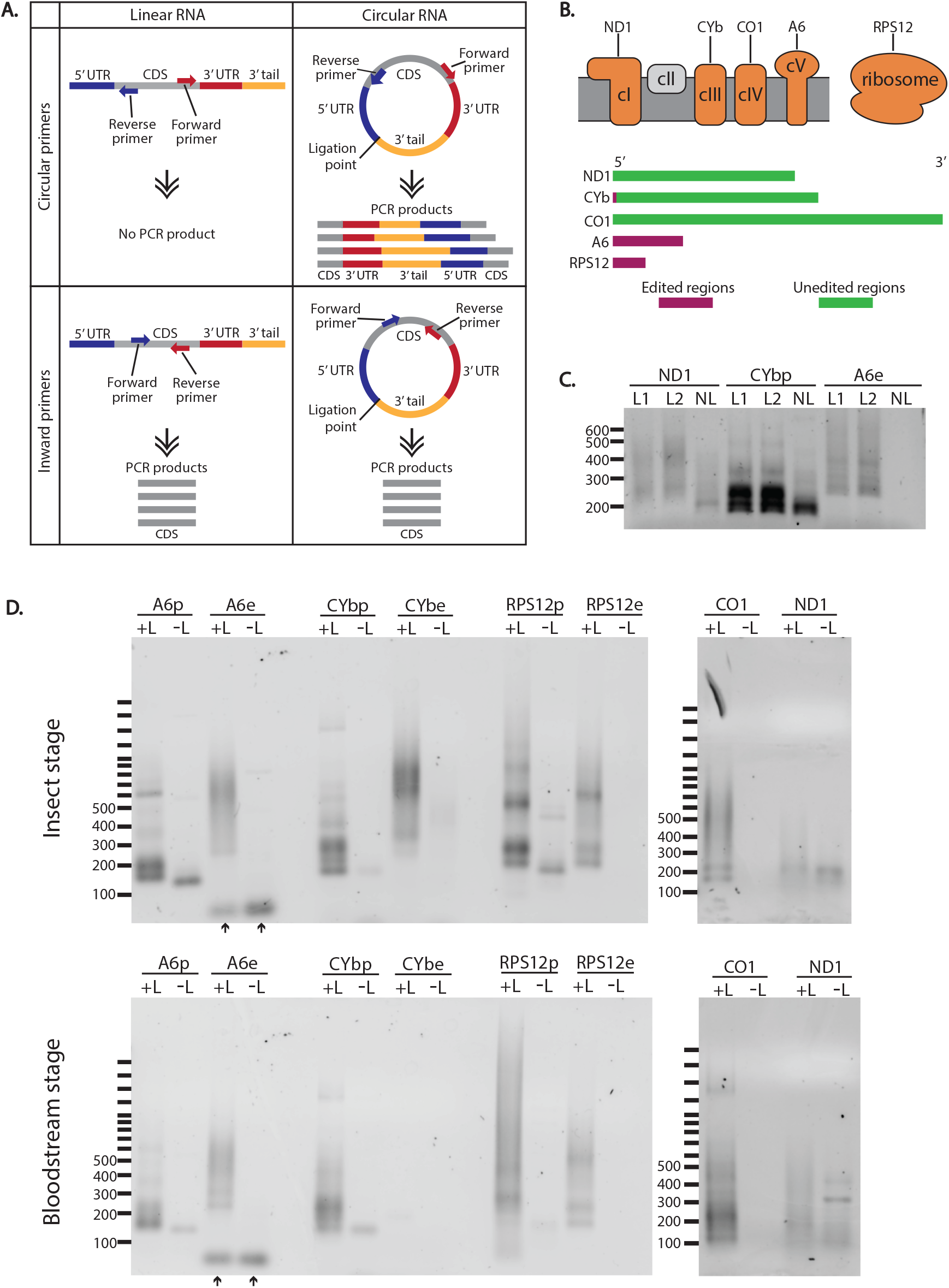
A subset of *Trypanosoma brucei* mitochondrial transcripts are circularized in the cell. Schematic of primer placement and PCR products from linear and circularized RNA. CDS, coding sequence. Schematic of the electron transport chain and mitochondrial ribosome with transcripts that were chosen for PCR analysis shown above the complexes of which they are a part (above). Complex II, shown in grey, does not have any subunits encoded in the mitochondrial genome. Length of bar is proportional to relative length of mRNA prior to editing (below). Green, regions that do not require editing prior to translation; maroon, regions that require editing prior to translation. *ND1*, NADH dehydrogenase subunit 1; *CYb*, cytochrome b; *CO1*, cytochrome c oxidase subunit 1; *A6*, ATP synthase subunit 6; *RPS12*, ribosomal protein S12. **(C)** Agarose gel of PCR products using circular oriented primers (top panels in A) for three different *T. brucei* mitochondrial transcripts from cDNA made from RNA of ‘control’ cells as defined in Materials and Methods. The smear of DNA is from the variation in tails lengths and is expected for these products. L1, T4 RNA ligase (Epicentre) added; L2, T4 RNA ligase (New England BioLabs) added; NL, no ligase added. Pre, pre-edited; e, edited. **(D)** Agarose gels of PCR products from RNA treated with ligase (+L) and without ligase (-L) of insect stage (top) and mammalian bloodstream stage (bottom) *T. brucei* for five mitochondrial transcripts. Transcripts that are edited have different primer sets to amplify their pre-edited (p) or edited (e) sequences. The blank well for +L *CYbe* in the bottom gel is expected because edited *CYb* transcripts are not found in bloodstream stage parasites.

We are currently using circTAIL-seq to investigate several mitochondrial transcript tail populations (**Figure 1B**) on newly generated parasite insect life stage cell lines that, when induced, will result in either overexpression and/or RNAi silencing of proteins of interest. Without the inducer tetracycline in the media, the cells are effectively wild type, lacking overexpression or silencing of any genes, thus gene expression mirrors that of the parent cell line 29-13 (referred to as ‘control cells’, and described further in Materials and Methods). In support of this, circTAIL-seq length and tail composition analysis from three of these cell lines in their uninduced state are practically indistinguishable **(Figure S1)**. In our current circTAIL-seq and in earlier experiments examining the effect of enzyme manipulation on tails by manually cloning and Sanger sequencing [18], the uninduced state of these cells are considered wild-type.

In generating amplicons for circTAIL-seq, we included a control PCR in which no ligase was added to the RNA, and thus should produce no amplicon library. Unexpectedly, we found that for some transcripts, PCR products were formed and visible by agarose gel electrophoresis even without the addition of ligase **(Figure 1C)**. The amplicon produced differed from those that were produced from *in vitro* ligated RNA in that products appeared less abundant, and the length of the products was shorter overall. These products are not byproducts of DNA contamination, as we failed to obtain PCR products with the same circular primers when purified DNA was used as a template (using the mitochondrial transcript *ND1*, **Figure S2A**). In contrast, an *ND1* product of similar size to that of **Figure 1C** was generated when DNase treated RNA was used to generate the cDNA in the upstream step **(Figure S2A)**. The most likely explanation for the existence of these PCR products generated in the absence of *in vitro* mRNA circularization is that a fraction of mitochondrial mRNAs is naturally and normally circularized in the parasite mitochondrion.

To determine the extent of such a phenomenon, we tested all our functional circular primer sets on cDNA of RNA from parent 29-13 insect life stage parasites and bloodstream-stage ‘single marker’ (SM) Lister 427 life stage parasites both with and without performing *in vitro* RNA circularization **(Table S1, Figure 1D)**. Normally, the circTAIL-seq workflow includes a cycle optimization step for each transcript (example shown in **Figure S2B)** in which we identify the number of cycles resulting in products of the expected size range with a minimal number of unusable higher molecular weight concatenated PCR products. However, *in vivo* circularized transcripts can be more clearly observed by gel electrophoresis with additional PCR cycles, so to best compare these amplicons to *in vitro* circularized amplicons, we increased cycles for all transcripts. Thus, some lanes in Figure 1D exhibit these high molecular weight artifacts as well as the expected products in the lower size range. For the insect life stage parasites, 4 of 5 tested transcripts produced products without ligase addition, and the products were shorter and qualitatively appeared less abundant than when ligase was added to the RNA. The result was the same for the bloodstream stage parasites.

Notably, for transcripts requiring editing, reliable products were detected for the pre-edited (p) but not the edited (e) versions of the tested mRNA. This suggests that if edited transcripts are circularized *in vivo*, their abundance is very low. Further, an amplicon was not generated for never-edited transcript *CO1* unless it was first circularized *in vitro*, indicating that whether a transcript undergoes editing may be related to is likelihood of being circularized. To confirm that some transcripts in the untreated samples did not produce a product with circular primers, we increased the amount of PCR cycles further (**Figure S2C)**. For insect stage RNA without ligase addition, products were now detected for *RPS12e, CYbe*, and *CO1. CO1* products were also detected in the untreated bloodstream form sample. But, given that higher cycles also produced a band for edited *CYb* for the ligase-treated bloodstream form RNA, which is known to be practically absent in this life stage of the parasites, the identity of these additional bands is unclear. We cloned and Sanger sequenced circular products from transcripts for NADH dehydrogenase subunit 1 (*ND1*), pre-edited cytochrome b (*CYbp*), and pre-edited ATP synthase subunit 6 (*A6p*) from insect life stage parasites. Sequencing confirmed that these products were indeed circRNA: 3’ ends of mRNAs, sometimes including a short tail sequence, covalently ligated to 5’ ends of the same transcript. We attempted to clone and sequence products from edited *CYb* (*CYbe*) but were unsuccessful.

### Circularized mRNA populations are distinct from total mRNA

With increased confidence that the amplicons generated were from native mitochondrial circRNA molecules, we further characterized them to distinguish how they may differ from linear mRNAs of the same transcript. We examined ligation point junction sequences between high-throughput sequencing reads from insect-stage parasite libraries obtained with and without *in vitro* circularization of RNA. We extracted the ligation point junction sequences from Illumina reads of libraries obtained from 29-13 strain parasites untreated RNA (sequences only from circRNAs) and *in vitro* circularized parasite RNA from control cells described above to identify any termini characteristics unique to circRNA (**Table S2)**. These latter read pools are believed to contain mostly linear mRNA reads due to the high amplicon abundance of these libraries relative to circRNA amplicons. However, as the linear to circularized mRNA ratios in parasites have not been quantified, we refer to these junction sequence populations as ‘total’ rather than ‘linear’. We examined libraries from three transcripts that range in their requirement for RNA editing. They were: never edited *ND1*; *CYb*, which is edited in only a small region; and *A6*, an extensively edited transcript **(Figure 1B)**. Since we were only able to verify true amplicons via Sanger sequencing using primers specific to the pre-edited forms of edited transcripts, we only examined *CYb* and *A6* transcript reads that had not yet entered the editing pathway (*CYb<u>p</u>* and *A6<u>p</u>*).

For each transcript, characteristics of the tail populations between ligation point junction sequences of total and circRNA libraries were compared first. It was clear that tails from circRNAs were shorter and contained far fewer As (**Figure 2)**. For instance, the highest proportion of *A6p* tails are between 10 and 15 nucleotides in length when the library derived from total RNA was examined, whereas the *A6p* derived from naturally circularized mRNAs generally incorporated tails of less than five nucleotides. Tail length trends for *CYbp* and *ND1* were similar (**Figure 2A**). Nucleotide compositional differences were also observed. For each transcript’s tail population, **Figure 2B** plots show the fraction of nucleotides that are A rather than U at each nucleotide position. For all three transcripts, tails of libraries derived from total mRNA possess regions where A content is as high as 75%. In contrast, tails derived from mRNA in its circular form have a dramatically lower A content, generally below 50% and maximally reaching 60% (**Figure 2B)**. To ensure that our characterizations of tail populations incorporated into circRNAs were not strain-dependent, we sequenced circRNA tail populations of the same transcripts in the EATRO164 Istar1 (EATRO164) cell line of a different lineage than that of the bloodstream and insect stage cells used thus far. Circularized mRNA tails from EATRO164 parasites were very similar to the circularized mRNA tails we found in 29-13 parasites **(Figure S3)**.

**Figure 2.**
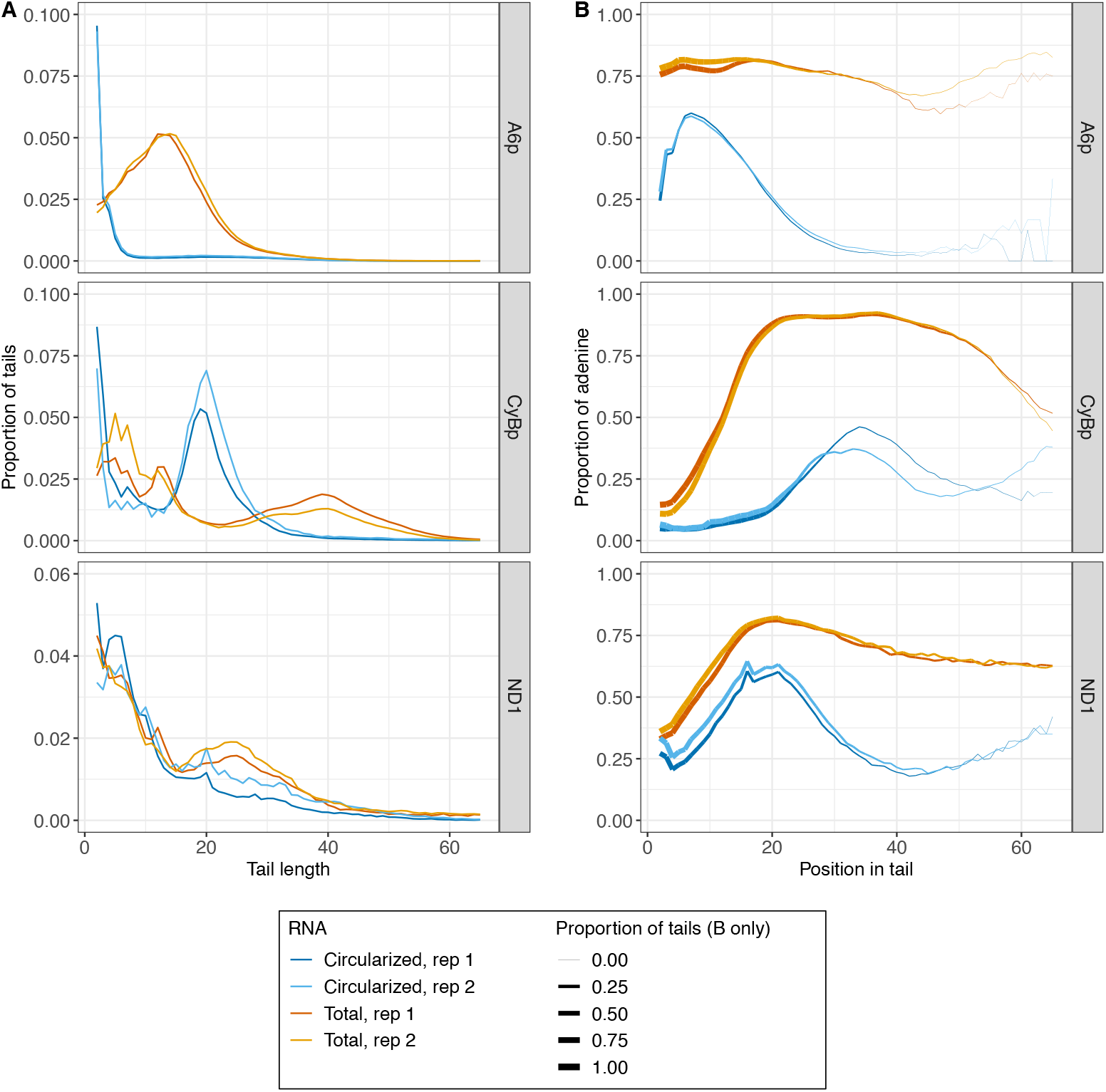
*Trypanosoma brucei* mitochondrial circularized mRNA (circRNA) possess shorter tails with lower adenine content than tails on the total mitochondrial mRNA of the same transcript. Populations of tails in circRNAs are shown in dark and light blue and tail populations from total mRNA are shown in dark and light orange. **(A)** Population density curves of tail lengths for three mitochondrial transcripts. **(B)** Population density curves of proportions of nucleotides that are adenine at each position along tails for three mitochondrial transcripts. The thickness of the line represents the proportion of tails that are long enough to contribute to the data at each nucleotide position. As the line becomes thinner, the proportion of tails that are available to contribute to the adenine content data approaches zero because there are very few long tails. Data is shown from the second nucleotide in the tail to account for ambiguity in tail cutoff determination.

Differences in length and composition between total RNA and circRNA tails suggest that the manner in which tails are being added to the mRNAs is distinct. Two enzymes add nucleotides to mRNA 3’ ends: KPAP1 adds As and KRET1 adds Us. To investigate how A and U addition may be different for circRNA and total RNA, we used hidden Markov modelling (HMM) [19]. To define the number of nucleotide addition ‘states’ required for HMM, the two types of mRNA tails on kinetoplastid mitochondrial mRNAs must be considered. The first is a shorter tail found on all transcripts that consists of infrequent switching between Us and As, which results in a succession of homopolymers of As (state 1) and/or Us (state 2). For never-edited transcripts such as *ND1*, these shorter tails are sometimes elongated with a sequence characterized by frequent switching between As and Us presumably involving both KPAP1 and KRET1 combined activities (state 3). Thus, although it is appropriate to model pre-edited transcripts *CYbp* and *A6p* with a 2-state model, a 3-state model is required for never-edited *ND1* (**Figure 3**). For *A6p* (**Figure 3, top row)** while the majority of tails in both the total and circularized RNA initially enter an A-adding state (shown in red), a higher percentage of circRNA enter the U-adding state (34-35%) than the total RNA (24-26%). Additionally, more circRNA tails move from an A-adding state to the U-adding state (14-15%) than total RNA (3%). This is reflected in the lower percentage of As in the circRNA tails shown in **Figure 2B**. Similar trends are shown for *CYbp* tails (**Figure 3, middle row)**.

**Figure 3.**
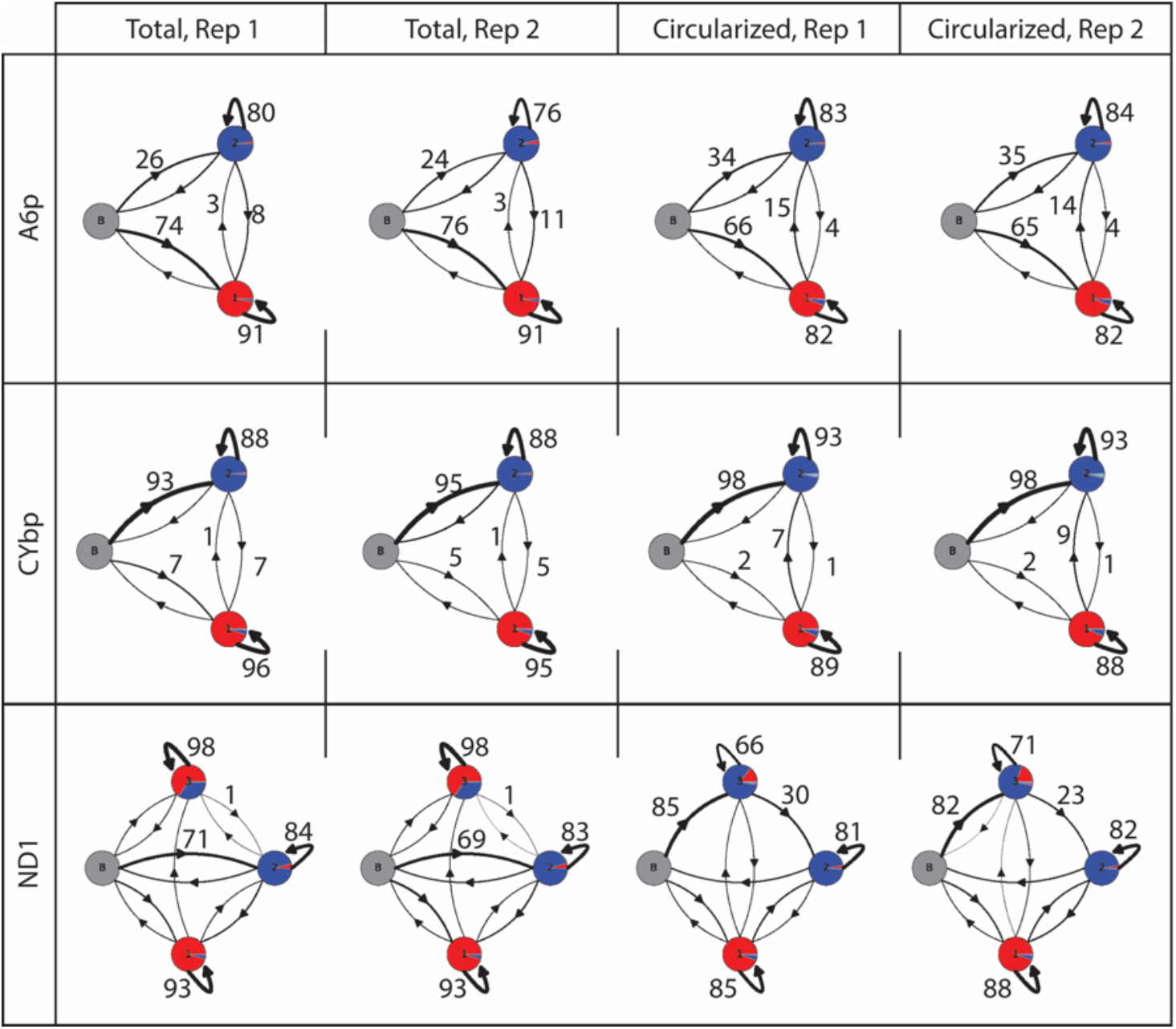
Hidden Markov modelling reveals differences in tail addition enzyme activities between total RNA and circRNA of the same transcript. Percentage of adenine addition (red) and uridine addition (blue) are shown for each state by proportion of the circle colored. The percentage of tails that move from one state to another is shown by thickness of arrows. Actual percentages are written next to arrows of particular interest. Grey, starting state; 1, primarily A-adding state; 2, primarily U-adding state; 3, A/U-adding state.

For *ND1* (**Figure 3, bottom row)**, while most tails of the total RNA population move into the primarily U-adding state immediately (69-71%), the majority of circRNA tails instead start in the A/U addition state (82-85%). Though this may first appear to contradict **Figure 2B**, there is a higher percentage of Us added in the A/U-adding state of circRNA when compared to total RNA (shown as relative proportions of red (A) and blue (U) in the circle). Further, many tails quickly exit this state of addition and proceed to the U-adding state on circRNA molecules (23-30%). In addition to confirming data seen in **Figure 2**, HMM gives us new insight into how the enzymes may be functioning differently during tail addition. The tails that stay in one state reflect processive enzyme activity, while the tails that move to another state suggest enzymes that are switching more frequently. In all three transcripts tested, the A-adding KPAP1 is less processive in circRNA tails than total RNA tails. This is shown by the decrease in the percentage of tails that stay in the A-adding state when comparing total RNA to circRNA (91% in total RNA to 82% in circRNA for *A6p*, 95-96% to 88-89% for *CYbp*, 93% to 85-88% for *ND1*). For the pre-edited transcripts (*A6p* and *CYbp*), we also see an increase in the percentage of tails that stay in the U-adding state (76-80% in total RNA to 83-84% in circRNA for *A6p*, 88% to 93% for *CYbp*), indicating that the U-adding KRET1 may be more processive. The HMM for *ND1* indicates a different enzymatic activity pattern, with the predominant tail percentage starting in the A/U-adding state, which indicates a less processive KRET1 that is often interrupted by KPAP1.

Considering that tails incorporated into circRNAs are shorter than those of the total mRNA population and that there are differences in tail-addition enzymatic activity between total and circRNAs, circularization may occur during mRNA processing around the step of tail addition initiation. If this is the case, we may expect a higher proportion of ligation point junctions in which the 3’ terminus entirely lacks an untemplated tail sequence in the circRNA libraries than in those of the total RNA-derived libraries. We determined the percentage of reads within ligation point junction regions that lack tail sequences, and found that for transcripts *ND1* and *A6p*, circRNAs indeed had a higher percentage of reads that lacked evidence of 3’ mRNA tail addition prior to circularization (**Figure 4A)**. Ligation point junction sequences from mitochondrial circRNAs of EATRO164 parasites revealed that similar percentages lacked tails (**Figure S4A)**. However, in both parasite cell lines *CYbp* circRNAs were tailed at percentages similar to that of total *CYbp* mRNAs. In the circRNA-derived libraries, most of the *CYbp* tails were short (< 25 nucleotides) oligo(U) sequences (**Figure 2)**. The tendency for *CYbp* tails to have short strings of U more predominantly at the beginning of tails compared to other transcripts has been noted previously [20]. This difference in tail composition suggests that *CYb* mRNA processing may differ from that of other mitochondrial transcripts. The fact that this difference was captured in the circRNA-derived sequence library suggests that differences in processing occur prior to *CYb* being vulnerable to covalent self-circularization.

**Figure 4.**
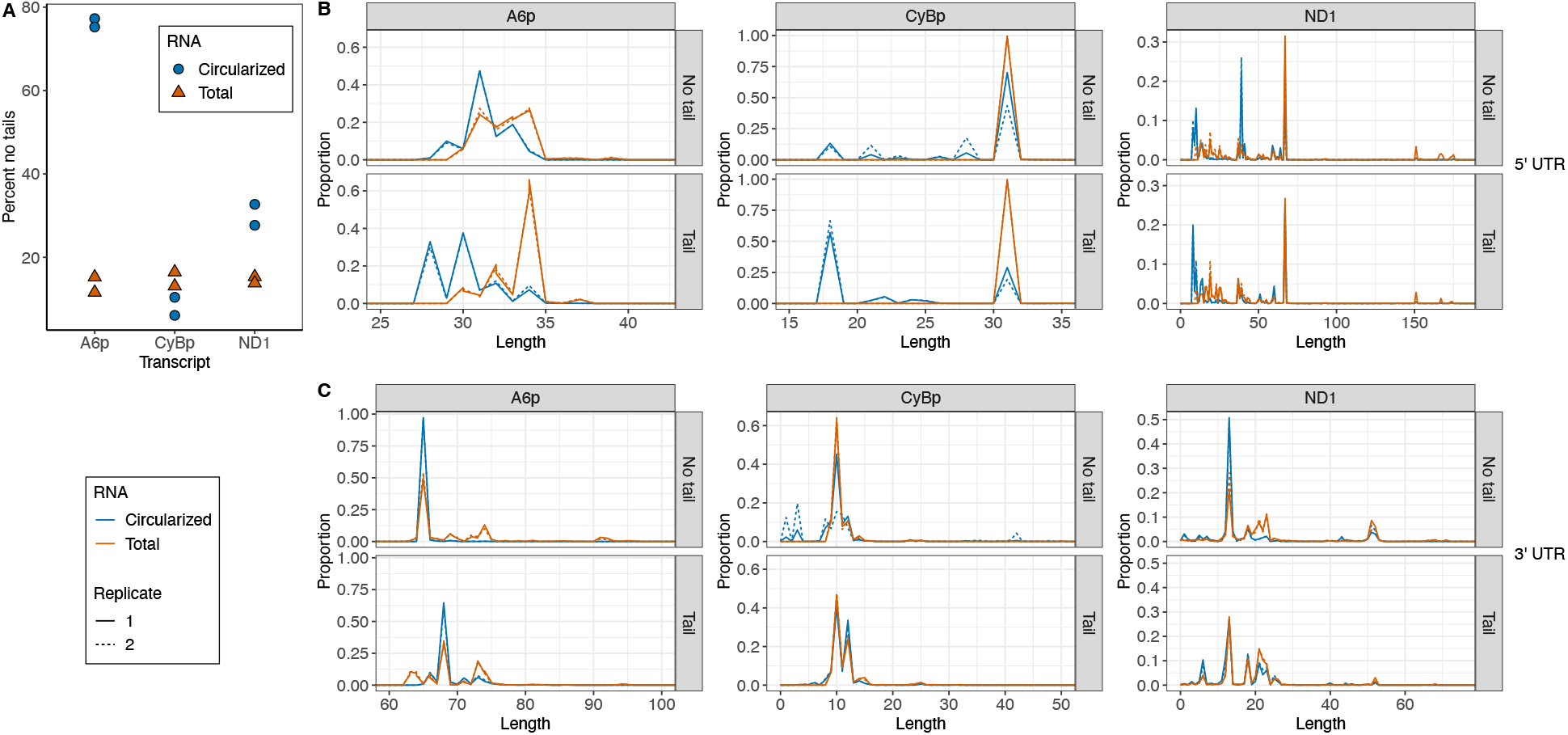
*Trypanosoma brucei* circularized mRNAs (circRNAs) have distinctly different 5’ termini and fewer tails than total mRNAs of the same transcript. **(A)** Percent of sequenced ligation point junction reads that possess no untemplated A or U additions between the ligated ends of the 3’ and 5’ UTRs. **(B)** 5’ UTR lengths of circRNA (blue) and total mRNA populations (orange) for each transcript. **(C)** 3’ UTR lengths of circRNA (blue) and total mRNA populations (orange) for each transcript. For (B) and (C), populations of 3’-5’ junctions that contain nontemplated tail sequence are analyzed separately from populations that do not. 5’ UTR lengths are counted starting with the first nucleotide after the tail ends until the first nucleotide of the start codon. 3’ UTR lengths are counted so that the last nucleotide after the stop codon is position +1. Replicate 1, solid line; replicate 2, dotted line.

Since tail addition on mitochondrial mRNAs that eventually became circularized was curtailed and altered, we considered that mRNAs vulnerable to circularization may display other types of altered maturation. 5’ and 3’ cleavage or trimming are also mitochondrial mRNA maturation steps, so we investigated the UTR lengths of circRNAs and total mRNAs. Using our high-throughput sequencing reads of ligation point junction regions, we determined the 5’ and 3’ UTR lengths between a transcript’s circularized mRNA and total mRNA population. As tail addition may be linked to proper UTR processing, we separately addressed the UTR length comparison of mRNAs that possess tail sequence and those that do not (**Figure 4, B and C)**. We found 5’ UTR lengths were shorter in a transcript’s circRNA population compared its total RNA population when there was no tail present (**Figure 4B, top panel**), and this difference increased when there was a tail present (**Figure 4B, bottom panel)**. However, we surprisingly found that the 3’ UTR length was consistent between a transcript’s circularized and total mRNA populations regardless of whether a tail was present (**Figure 4C)**. UTRs of circRNAs from the EATRO164 cell line were comparable in length to those of strain 29-13 (**Figure S4 B and C)**, with the exception of some differences in 3’ UTRs of *A6p* that may be related to sequence differences between the strains.

Considering our discovery that circRNA ligated termini have characteristics that differ from termini of linear mRNAs, it was important to verify the robustness of our sequence dataset from those libraries. We recognized that high-throughput approaches utilizing PCR for library generation can suffer from library low diversity, resulting in less information content in the sequencing output than read numbers would suggest. We previously demonstrated that libraries of traditional circTAIL-seq mitochondrial transcripts employing *in vitro* circularization of linear mitochondrial transcripts from *T. brucei* were sufficiently diverse [17]. Knowing that starting template numbers of circularized molecules used to generate circRNA libraries are likely low, we examined our circRNA library read pools for sequence diversity. First, we looked at the incidence of reads that contained identical junction regions (which include all sequence between the primer annealing positions) (**Table S2)**. The sequences appearing the most in each library that aligned with the DNA template were always 3’-5’ termini direct ligations lacking tail sequence or 5’ and 3’ ends interspersed with a short mono-nucleotide tail sequence. This is not surprising because our sequencing captures a snapshot of the processing state of the population of a transcript. All molecules on their way to maturity will have to pass through a state of no tail or short oligo tail. Further, in reads lacking tails or possessing only short oligonucleotides, the lengths of 3’ and 5’ UTRs did vary within the range shown in **Figure 4**, indicating that unique molecules were being represented in the read population. We did not see highly duplicated reads with tails that had longer, unique A/U patterns. The repeated appearance of such a sequence among the reads would suggest a single template molecule amplified repeatedly. Additionally, we identified the quartiles for each library for all reads (**Table S2)** and found that while there are highly repeated sequences as identified above, most junction regions were duplicated only 2 to 9 times. We conclude that while the total number of sequences analyzed in our circRNA libraries were typically lower than those of the total RNA libraries (**Table S2**), these populations appear to be sufficiently diverse for the analyses that we performed.

In summary, when compared to a transcript’s total mRNA population, its circRNA population exhibits shortened 5’ UTRs and altered non-templated tail characteristics, but consistent 3’ UTR termini. Together, these data support an mRNA processing model in which 1) modification of the 3’ and 5’ ends of mRNA are physically linked, and 2) circularization happens after the MPsome has trimmed the 3’ UTR and added an initial oligo(U) tail, but before the oligo(U) tails acquire A-rich extensions.

## Discussion

We have shown for the first time that there are mitochondrial-encoded circular mRNAs in kinetoplastids. We detected that circRNAs likely comprise part of the total mRNA population for 4 of the 5 transcripts tested in both the insect-stage and bloodstream stage parasites, though for transcripts requiring editing, only the pre-edited versions showed convincing evidence of circularization (**Figure 1D**). After high-throughput sequencing three transcripts in two different strains, we confirmed that trypanosomatid mitochondrial circRNAs are not isolated to one strain of *T. brucei* and that mitochondrial circRNA in the two strains have similar characteristics (**Figures S3 and S4**). We determined that these circRNAs are a subpopulation of mitochondrial mRNAs that differ in their 3’ non-encoded tail and 5’ UTR characteristics relative to their linear counterparts (**Figures 2 and 4**), and that enzymatic activity on circRNAs during tail addition is different, in that it is more limited, than the activities acting on total RNA (**Figure 3**). These population-level differences strongly suggest that the sequenced ligation point junctions of circularized molecules are not an artifact of the amplification and sequencing protocols used but rather come from molecules naturally existing in *T. brucei* mitochondria. The presence of circRNA suggests a new pathway in the *T. brucei* mitochondrial mRNA lifecycle.

Since a primary mechanism of trypanosome mitochondrial mRNA processing has been proposed and proteins involved identified, we can derive models for how and when circRNAs are formed. We present one model in **Figure 5**. Processing complexes on mRNA 5’ and 3’ ends are thought to bring the mRNA termini near to each other through protein-protein interactions between the complexes [13]. 5’ and 3’ end proximity would be advantageous for subsequent mRNA covalent circularization.

**Figure 5.**
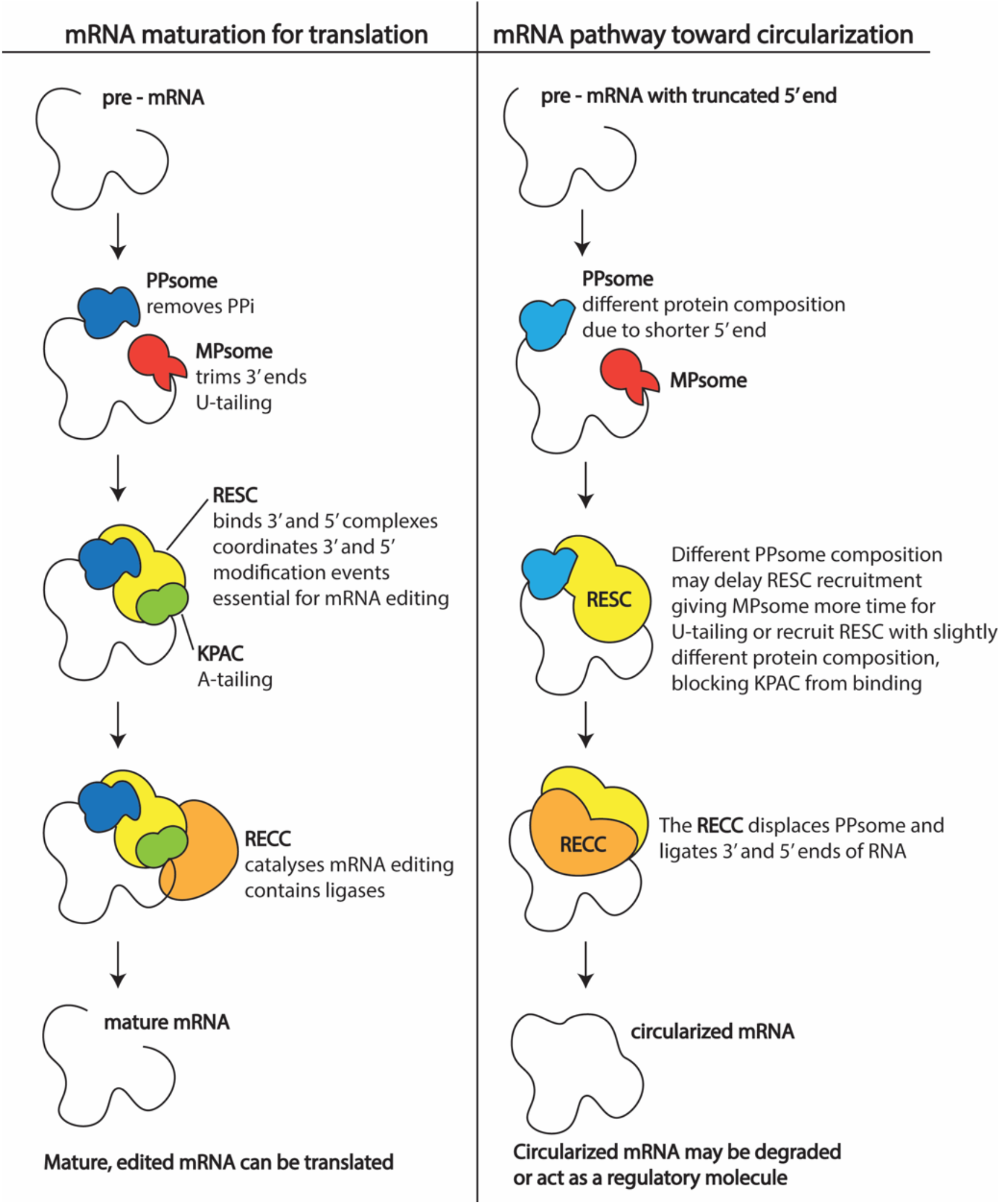
Model of potential mRNA maturation pathways for translatable transcripts (left) and circularized transcripts (right). PPsome, 5’ pyrophosphohydrolase complex; MPsome, mitochondrial 3’ processome; RESC, RNA editing substrate-binding complex; KPAC, kinetoplast polyadenylation complex; RECC, RNA editing catalytic complex.

Our model suggests that protein complexes bring mRNA ends close to each other during processing earlier than what is currently believed. Interactions between proteins bound to 5’ and 3’ ends are postulated to occur after the addition of the short A-rich tail [13], rendering the mRNA resistant to further trimming and oligo U-tailing. Sequenced circRNAs have similar 3’ UTR lengths as mRNAs of the total transcript population, yet fewer non-encoded 3’ nucleotide additions (**Figure 4)**. CircRNA that are tailed typically have U tails rather than the A-rich tails characteristic of linear mRNA (**Figure 2)**. This suggests that protein interactions between the 3’ and 5’ bound complexes bring termini together before global KPAP1 A-tailing of mRNAs but after MPsome trimming and U-tailing. Thus, A-rich nucleotide 3’ extensions may not be necessary to promote a protective mRNA configuration.

Factors capable of driving processing toward covalent circularization as opposed to the normal steps of RNA editing and translation include characteristics of the transcript itself. For instance, the shortened or altered length of 5’ UTRs of circRNAs could favor covalent circularization. The observed 5’ UTR variability itself could arise from exoribonuclease or endoribonuclease activity, or alternative transcription start sites. Regardless of how they arise, 5’ UTR length differences could influence the composition of 5’ UTR-bound proteins (**Figure 5**). Such differences could serve to delay A-rich tail addition on the 3’ end, leaving the mRNA vulnerable to covalent circularization by ligases.

Two ligases function in editing, and editing machinery interacts with 5’ and 3’ complexes of all transcripts, not just those that are edited [14]. Thus, it is possible that the catalytic RECC that contains these ligases may be positioned such that it can circularize transcripts with variable 5’ UTRs. Further evidence that mRNA circularization may be carried out by editing ligases is that the pre-edited transcripts *CYb* and *A6*, which would need to interact substantially with several editing complexes to initiate editing, seem to have more circRNAs than the never edited *ND1*. This is shown in the percentage of reads that aligned to the respective transcripts (**Table S2**). Only a fraction of the material sequenced in the *ND1* circRNA libraries aligned to the 3’ to 5’ DNA template, suggesting that these circRNAs are less abundant for never-edited transcripts. Although beyond the scope of this study, quantitatively testing this hypothesis using RT-qPCR would be ideal and designing these experiments is planned for future work.

Even though mitochondrial mRNA circularization may involve factors and enzymes normally thought to execute other functions, it does not mean that it is unregulated or that it does not have a function. Possible functional roles for mitochondrial cirRNAs include a role in mRNA degradation. Atypical 5’ UTR lengths could indicate incorrect transcription initiation or an otherwise compromised mRNA, and circularization may mark such mRNA for degradation. Various methods for circRNA degradation have been proposed [21], though some circRNAs have been shown to be more stable than their linear counterparts [22,23]. If *T. brucei* mitochondrial circ RNAs are stable rather than unstable, they could serve as regulators. In other systems circRNAs regulate various processes by acting as microRNA sponges [24], affecting mitochondrial protein import [11], and regulating translation or transcription of certain proteins [25]. Though recently reported circularized mitochondrial mRNA have been shown to be translated in plants [10], in *T. brucei* this is unlikely because we primarily observe circRNAs formed from pre-edited transcripts that have not yet acquired an open reading frame.

Additionally, since circularization seems to halt proper mRNA tailing and is associated with shorter and more variable 5’ UTRs, this suggests circRNAs are not later translated, but are rather directed into a new RNA pathway entirely.

In this study, we show that a subset of *T. brucei* mitochondrial mRNAs are circularized early in the mRNA maturation lifecycle. The mechanism of circularization and potential role of circRNA in the mitochondrial mRNA lifecycle is intriguing and furthers our understanding of the temporal steps of trypanosome mRNA maturation.

## Materials and methods

### Parasite cell culture and cell lines

*Trypanosoma brucei* 29-13 cells (Lister 427 strain expressing T7 polymerase and tetracycline repressor), and EATRO164 Istar1 (EATRO164) insect-stage cells were grown in SDM-79 at 27°C in 5% CO2. SDM-79 was supplemented with G418 and hygromycin for 29-13 cell growth. Bloodstream-stage ‘single marker’ (SM) Lister 427 *T. brucei* cells were grown at 37°C in 5% CO2 in HMI-9 supplemented with G418. Most experiments to detect and characterize mitochondrial circRNAs, including all deep sequencing of circRNAs, were performed on 29-13, EATRO164, and SM parasite RNA. However, mitochondrial circRNAs were first identified with gel electrophoresis and Sanger sequencing on 29-13 parasites that have an additional nonphenotypic genetic modification. Currently unpublished and termed “control” here, the cell line is identical to 29-13 except that, under induction with tetracycline, it would express the protein KPAP1 from an exogenous locus and simultaneously silence its endogenous KPAP1 protein. Lacking tetracycline in its growth medium, this cell line is phenotypically identical to 29-13 in growth and other parameters. circTAIL-seq (including an *in vitro* ligation step) of total transcript mRNA (which is likely majority non-circular in nature) was performed on this cell line. These same data from “control” serve as wild-type control in a different study to be published elsewhere. “Control” cell line medium was supplemented with G418, hygromycin, puromycin, and phleomycin.

### RNA manipulations and PCR product gel electrophoresis analysis

RNA was collected as described [26]. RNA was circularized using T4 RNA ligase 1 (Epicentre or New England BioLabs) and reverse transcribed using Superscript III (Invitrogen) and gene specific primers as in [17] with all reverse transcription primers listed in **Table S1**. ‘No ligase’ control samples were obtained by substituting water for ligase in the reaction. For PCR utilizing circular primers (**listed in Table S1)**, product was generated with KAPA2G Robust polymerase (KAPA biosystems), manufacturer provided buffer per manufacturer’s protocol. Variable amounts of cycles and cDNA were used because the concentration of cDNA varied greatly depending on the preceding manipulations: a 1:10 dilution of cDNA and 33 cycles was used for **Figure 1C**; a 1:4 dilution of cDNA was used in PCRs of ligase and mock ligase treated samples using 30 cycles in **Figure 1D** and 35 cycles in **Figure S2C**; and undiluted cDNA and 28 cycles was used for **Figure S2A**. For PCR conditions, after an initial incubation at 95°C for 3 minutes, cycling was performed as stated as follows: 95°C for 0:15, annealing temperature listed in **Table S1** 0:15, and extension at 72°C for 0:30. A final extension step at 72°C for 1 minute was performed. All reactions were electrophoresed on 1.5% ultrapure agarose gels.

### Cloning and Sanger sequencing

50µl PCR products generated from no-ligase-added cDNA primed with circular primers for *ND1, CYbp*, and *A6p* (**Table S1**) were run on agarose gels and purified using Freeze ‘N Squeeze DNA gel extraction spin columns (Bio-Rad). For manual sequencing, purified PCR products were cloned using NEB PCR Cloning Kit (New England BioLabs). Clones were Sanger sequenced at the University of Minnesota Genomics Center.

### CircTAIL-seq

CircTAIL-seq was performed as described in [17] with modifications to PCR cycling conditions, sequencing read lengths obtained, and parts of downstream read processing. Reverse transcription primers, Illumina primers, and annealing temperatures are listed in **Table S1**. For cDNA generated from RNA treated with ligase, a 1:10 dilution of cDNA was used in PCRs, as per the circTAIL-seq protocol in [17], and PCR cycling conditions were specific for each primer set (**Table S1)** to decrease the amount of concatenated high molecular weight products as is demonstrated in **Figure S2B** for *A6p*. For cDNA generated from RNA not treated with ligase, cDNA was not diluted prior to PCR, and PCRs were cycled 34 times to increase the abundance of the products to be sequenced. The University of Minnesota Genomics Center (UMGC) performed quality control on all libraries by assessing quality and quantity on an Agilent BioAnalyzer and performing KapaQC. Amplicon libraries were then sequenced at UMGC on an Illumina MiSeq analyzer using the MiSeq V2 Chemistry 150 PE kit, acquiring 150bp paired-end reads. One exception, CyBp control replicate 1, was sequenced using MiSeq V2 Chemistry 250 PE kit. Eight libraries were run in one lane and sorted using barcodes when possible and unique gene primer sequences otherwise. **Table S2** lists the numbers of raw reads for each sequence file and, following processing, the number of merged reads utilized for downstream analysis of tails and UTR lengths. Though between 96-99% of reads were maintained after merging, removing reads that did not align to UTR templates and those that were partially edited resulted in 72-91% of reads used in downstream analysis for most libraries. The exceptions were the ND1 libraries, in which only 51-57% of reads were used for the control libraries and between 8-19% were used in the circularized libraries. Initial downstream read processing was performed as in [17].

Briefly, read quality was assessed with FastQC, paired reads were merged using PEAR, Illumina barcodes were trimmed with Trimmomatic, reads were reverse complemented and converted to fasta files using the FastX toolkit, and merged read quality was checked using FastQC. An updated tail identification protocol was used to identify which nucleotides constituted tail sequence for each read and which were part of the 3’ and 5’ UTRs surrounding any tail. The Needleman Wunch algorithm [27], coded in C, was modified for the alignments, while all downstream analysis was done by Python scripts (all code available at https://github.com/Pooryamb/circTAIL). Reads within files were aligned to the DNA sequence corresponding to both the 3’ sequence immediately downstream of the primer binding site to identify the last 3’ templated nucleotide for each read and to the 5’ region just upstream of the 5’ primer binding site to identify the 5’ last untemplated nucleotide. These alignments returned files of 3’ ends, 5’ ends, and non-encoded tails. Because the aligner predicts the longest possible UTR, tails were occasionally mis-aligned to A/U rich UTRs. To address this, we implemented a pruning step in circTAIL-seq alignment. Pruning starts from the end of the alignment and, based on a transcript-specific scoring matrix, terminates the 3’ and 5’ UTR alignment at the point where an A/U rich region follows a gap or mismatch that cannot be attributed to RNA insertion/deletion editing. For EATRO164 *A6p*, the DNA template used for alignment was modified to accommodate differences from the 29-13 *A6p* DNA sequence in the 5’ terminus.

### Hidden Markov Modelling

Hidden Markov modelling (HMM) was performed as described in [28] and [19].

## Acknowledgements

This work was supported by National Institutes of Health 1R15AI135885-01 awarded to SLZ, a research grant, 252733, from CIHR to RS, and the Robert Harpur foundation to PMB. CS was supported by funding from the University of Minnesota Graduate School Frieda M. Kunze Fellowship. The authors acknowledge the University of Minnesota Genomics Center (UMGC) for providing resources that contributed to the research results reported in this paper. The authors would like to thank Drs. Jason Smoniewski and Daniel Levings for assistance with Bash and R scripts used in data analysis.

## References

1. Sanger, H.L.; Klotz, G.; Riesner, D.; Gross, H.J.; Kleinschmidt, A.K. Viroids Are Single-Stranded Covalently Closed Circular RNA Molecules Existing as Highly Base-Paired Rod-like Structures. Proc Natl Acad Sci U S A 1976, 73, 3852–3856.

2. Hsu, M.-T.; Coca-Prados, M. Electron Microscopic Evidence for the Circular Form of RNA in the Cytoplasm of Eukaryotic Cells. Nature 1979, 280, 339–340, doi:10.1038/280339a0.

3. Arnberg, A.C.; Van Ommen, G.-J.B.; Grivell, L.A.; Van Bruggen, E.F.J.; Borst, P. Some Yeast Mitochondrial RNAs Are Circular. Cell 1980, 19, 313–319, doi:10.1016/0092-8674(80)90505-X.

4. Wang, P.L.; Bao, Y.; Yee, M.-C.; Barrett, S.P.; Hogan, G.J.; Olsen, M.N.; Dinneny, J.R.; Brown, P.O.; Salzman, J. Circular RNA Is Expressed across the Eukaryotic Tree of Life. PLOS ONE 2014, 9, e90859, doi:10.1371/journal.pone.0090859.

5. Danan, M.; Schwartz, S.; Edelheit, S.; Sorek, R. Transcriptome-Wide Discovery of Circular RNAs in Archaea. Nucleic Acids Res 2012, 40, 3131–3142, doi:10.1093/nar/gkr1009.

6. Salzman, J.; Gawad, C.; Wang, P.L.; Lacayo, N.; Brown, P.O. Circular RNAs Are the Predominant Transcript Isoform from Hundreds of Human Genes in Diverse Cell Types. PLoS One 2012, 7, e30733, doi:10.1371/journal.pone.0030733.

7. Lasda, E.; Parker, R. Circular RNAs: Diversity of Form and Function. RNA 2014, 20, 1829–1842, doi:10.1261/rna.047126.114.

8. Cahoon, A.B.; Qureshi, A.A. Leaderless MRNAs Are Circularized in Chlamydomonas Reinhardtii Mitochondria. Curr Genet 2018, 64, 1321–1333, doi:10.1007/s00294-018-0848-2.

9. Mance, L.G.; Mawla, I.; Shell, S.M.; Cahoon, A.B. Mitochondrial MRNA Fragments Are Circularized in a Human HEK Cell Line. Mitochondrion 2020, 51, 1–6, doi:10.1016/j.mito.2019.11.002.

10. Liao, X.; Li, X.-J.; Zheng, G.-T.; Chang, F.-R.; Fang, L.; Yu, H.; Huang, J.; Zhang, Y.-F. Mitochondrion-Encoded Circular RNAs Are Widespread and Translatable in Plants. Plant Physiology 2022, 189, 1482–1500, doi:10.1093/plphys/kiac143.

11. Zhao, Q.; Liu, J.; Deng, H.; Ma, R.; Liao, J.-Y.; Liang, H.; Hu, J.; Li, J.; Guo, Z.; Cai, J.; et al. Targeting Mitochondria-Located CircRNA SCAR Alleviates NASH via Reducing MROS Output. Cell 2020, 183, 76-93.e22, doi:10.1016/j.cell.2020.08.009.

12. Liu, X.; Wang, X.; Li, J.; Hu, S.; Deng, Y.; Yin, H.; Bao, X.; Zhang, Q.C.; Wang, G.; Wang, B.; et al. Identification of MecciRNAs and Their Roles in the Mitochondrial Entry of Proteins. Sci. China Life Sci. 2020, 63, 1429–1449, doi:10.1007/s11427-020-1631-9.

13. Aphasizheva, I.; Alfonzo, J.; Carnes, J.; Cestari, I.; Cruz-Reyes, J.; Göringer, H.U.; Hajduk, S.; Lukeš, J.; Madison-Antenucci, S.; Maslov, D.A.; et al. Lexis and Grammar of Mitochondrial RNA Processing in Trypanosomes. Trends in Parasitology 2020, 36, 337–355, doi:10.1016/j.pt.2020.01.006.

14. Sement, F.M.; Suematsu, T.; Zhang, L.; Yu, T.; Huang, L.; Aphasizheva, I.; Aphasizhev, R. Transcription Initiation Defines Kinetoplast RNA Boundaries. PNAS 2018, 115, E10323–E10332, doi:10.1073/pnas.1808981115.

15. Read, L.K.; Myler, P.J.; Stuart, K. Extensive Editing of Both Processed and Preprocessed Maxicircle CR6 Transcripts in Trypanosoma Brucei. Journal of Biological Chemistry 1992, 267, 1123–1128, doi:10.1016/S0021-9258(18)48405-0.

16. Suematsu, T.; Zhang, L.; Aphasizheva, I.; Monti, S.; Huang, L.; Wang, Q.; Costello, C.E.; Aphasizhev, R. Antisense Transcripts Delimit Exonucleolytic Activity of the Mitochondrial 3′ Processome to Generate Guide RNAs. Molecular Cell 2016, 61, 364–378, doi:10.1016/j.molcel.2016.01.004.

17. Gazestani, V.H.; Hampton, M.; Abrahante, J.E.; Salavati, R.; Zimmer, S.L. CircTAIL-Seq, a Targeted Method for Deep Analysis of RNA 3′ Tails, Reveals Transcript-Specific Differences by Multiple Metrics. RNA 2016, 22, 477–486, doi:10.1261/rna.054494.115.

18. Zimmer, S.L.; McEvoy, S.M.; Menon, S.; Read, L.K. Additive and Transcript-Specific Effects of KPAP1 and TbRND Activities on 3′ Non-Encoded Tail Characteristics and MRNA Stability in Trypanosoma Brucei. PLOS ONE 2012, 7, e37639, doi:10.1371/journal.pone.0037639.

19. Hampton, M.; Galey, M.; Smoniewski, C.; Zimmer, S.L. Probabilistic Models of Biological Enzymatic Polymerization. PLOS ONE 2021, 16, e0244858, doi:10.1371/journal.pone.0244858.

20. Kao, C.-Y.; Read, L.K. Targeted Depletion of a Mitochondrial Nucleotidyltransferase Suggests the Presence of Multiple Enzymes That Polymerize MRNA 3′ Tails in Trypanosoma Brucei Mitochondria. Mol Biochem Parasitol 2007, 154, 158–169, doi:10.1016/j.molbiopara.2007.04.014.

21. Zhou, M.; Xiao, M.-S.; Li, Z.; Huang, C. New Progresses of Circular RNA Biology: From Nuclear Export to Degradation. RNA Biol 18, 1365–1373, doi:10.1080/15476286.2020.1853977.

22. Jeck, W.R.; Sorrentino, J.A.; Wang, K.; Slevin, M.K.; Burd, C.E.; Liu, J.; Marzluff, W.F.; Sharpless, N.E. Circular RNAs Are Abundant, Conserved, and Associated with ALU Repeats. RNA 2013, 19, 141, doi:10.1261/rna.035667.112.

23. Enuka, Y.; Lauriola, M.; Feldman, M.E.; Sas-Chen, A.; Ulitsky, I.; Yarden, Y. Circular RNAs Are Long-Lived and Display Only Minimal Early Alterations in Response to a Growth Factor. Nucleic Acids Res 2016, 44, 1370–1383, doi:10.1093/nar/gkv1367.

24. Hansen, T.B.; Jensen, T.I.; Clausen, B.H.; Bramsen, J.B.; Finsen, B.; Damgaard, C.K.; Kjems, J. Natural RNA Circles Function as Efficient MicroRNA Sponges. Nature 2013, 495, 384–388, doi:10.1038/nature11993.

25. Li, Z.; Huang, C.; Bao, C.; Chen, L.; Lin, M.; Wang, X.; Zhong, G.; Yu, B.; Hu, W.; Dai, L.; et al. Exon-Intron Circular RNAs Regulate Transcription in the Nucleus. Nat Struct Mol Biol 2015, 22, 256–264, doi:10.1038/nsmb.2959.

26. Shaw, A.K.; Kalem, M.C.; Zimmer, S.L. Mitochondrial Gene Expression Is Responsive to Starvation Stress and Developmental Transition in Trypanosoma Cruzi. mSphere 2016, 1, e00051–16, doi:10.1128/mSphere.00051-16.

27. Eddy, S.R. What Is Dynamic Programming? Nat Biotechnol 2004, 22, 909–910, doi:10.1038/nbt0704-909.

28. Gazestani, V.H.; Hampton, M.; Shaw, A.K.; Salavati, R.; Zimmer, S.L. Tail Characteristics of Trypanosoma Brucei Mitochondrial Transcripts Are Developmentally Altered in a Transcript-Specific Manner. International Journal for Parasitology 2018, 48, 179–189, doi:10.1016/j.ijpara.2017.08.012.

29. Mesitov, M.V.; Yu, T.; Suematsu, T.; Sement, F.M.; Zhang, L.; Yu, C.; Huang, L.; Aphasizheva, I. Pentatricopeptide Repeat Poly(A) Binding Protein KPAF4 Stabilizes Mitochondrial MRNAs in Trypanosoma Brucei. Nature Communications 2019, 10, 146, doi:10.1038/s41467-018-08137-2.

30. Etheridge, R.D.; Aphasizheva, I.; Gershon, P.D.; Aphasizhev, R. 3′ Adenylation Determines MRNA Abundance and Monitors Completion of RNA Editing in T. Brucei Mitochondria. EMBO J 2008, 27, 1596–1608, doi:10.1038/emboj.2008.87.

31. Carnes, J.; Trotter, J.R.; Ernst, N.L.; Steinberg, A.; Stuart, K. An Essential RNase III Insertion Editing Endonuclease in Trypanosoma Brucei. Proc Natl Acad Sci U S A 2005, 102, 16614–16619, doi:10.1073/pnas.0506133102.

