## Supplemental Figures for "Circular mitochondrial-encoded mRNAs are a distinct subpopulation of mitochondrial mRNA in *Trypanosoma brucei*"

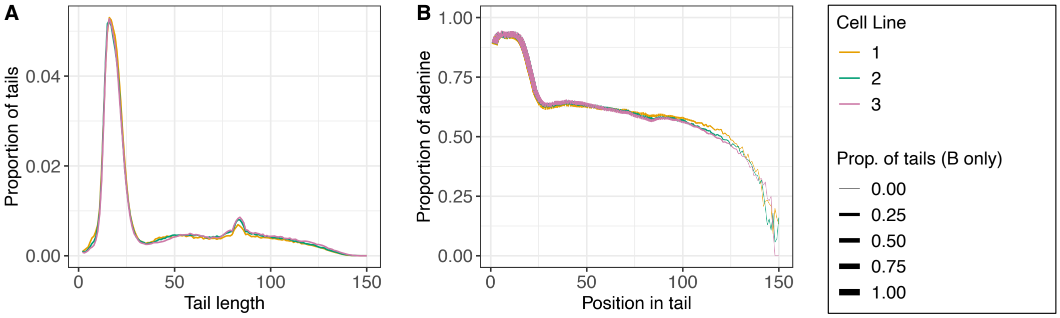


**Figure S1.** Three different uninduced control cell lines have similar tail characteristics for *Trypanosoma brucei* mitochondrial transcript *CO1*. **(A)** Tail length population density curves for the *CO1* transcript. **(B)** Population density curves of aggregate proportions of nucleotides that are adenine at each position along each tail for the *CO1* transcript. The thickness of the line represents the proportion the tail population that is long enough to contribute to the data at each nucleotide position. As lines becomes thinner, adenine content data is supported by fewer total tails because very few long tails are represented in the populations.

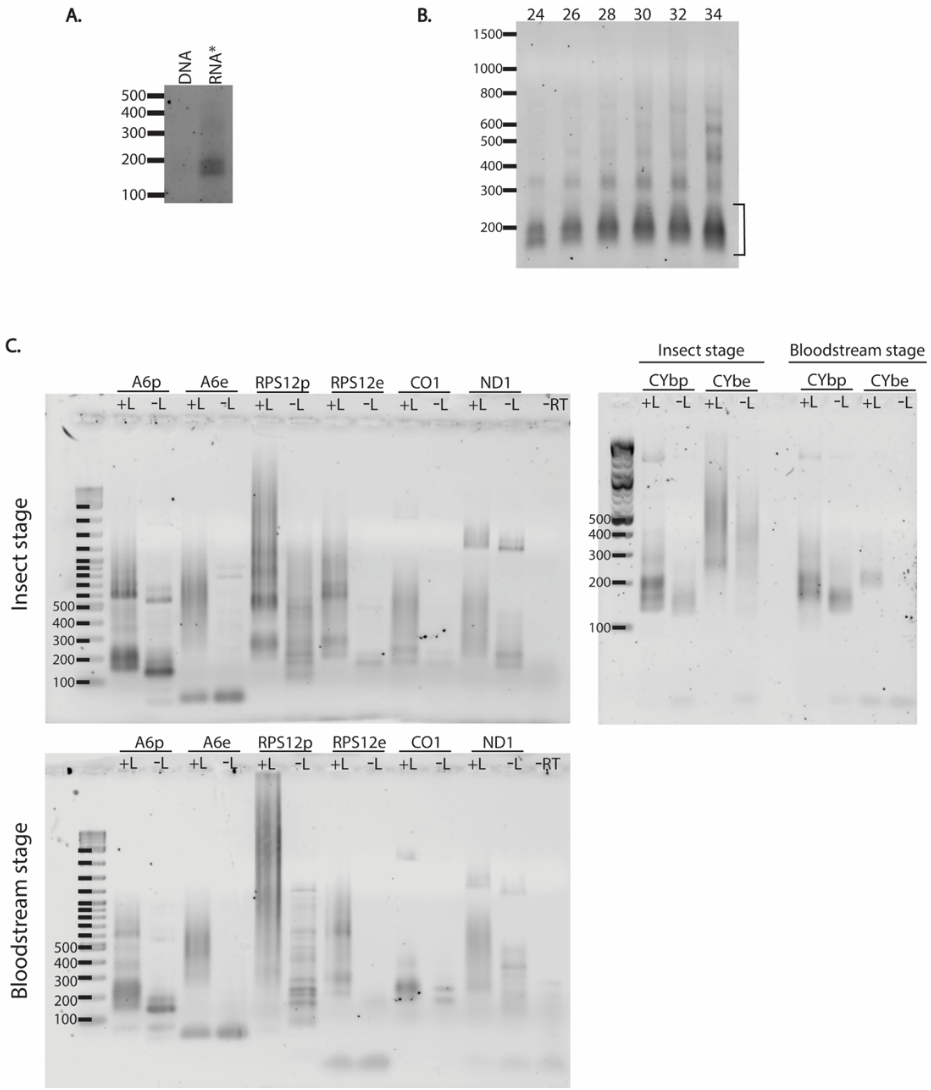


**Figure S2.** PCR evidence suggests that a fraction of mRNA in *Trypanosoma brucei* is circularized. **(A)** PCR products from circular primers are not the result of DNA contamination. Agarose gel electrophoresis of *ND1*

amplicon utilizing circular primers in PCR reactions. Lanes: DNA, 22ng of genomic DNA used directly as a PCR template; RNA*, reaction performed as described in Materials and Methods using DNase treated RNA to generate the cDNA as a template for the PCR reaction. **(B)** CircTAIL-seq PCR cycle optimization for *A6p* demonstrates that an optimal cycle range exists to obtain amplicons containing ligation point junction sequences without generating high molecular weight concatenated products that result from further amplification of the original low molecular weight products. Cycle numbers are shown at top of gel. Bracket indicates the desired product range. **(C)** Agarose gels of products of 35 cycle PCR reactions using as a template cDNA generated from RNA treated with ligase (+L) and without ligase (-L) of insect stage (top) and mammalian bloodstream stage (bottom) *T. brucei* for five mitochondrial transcripts. Transcripts that are edited utilize different primer sets to detect their pre-edited (p) and edited (e) versions. -RT, minus reverse transcriptase control.

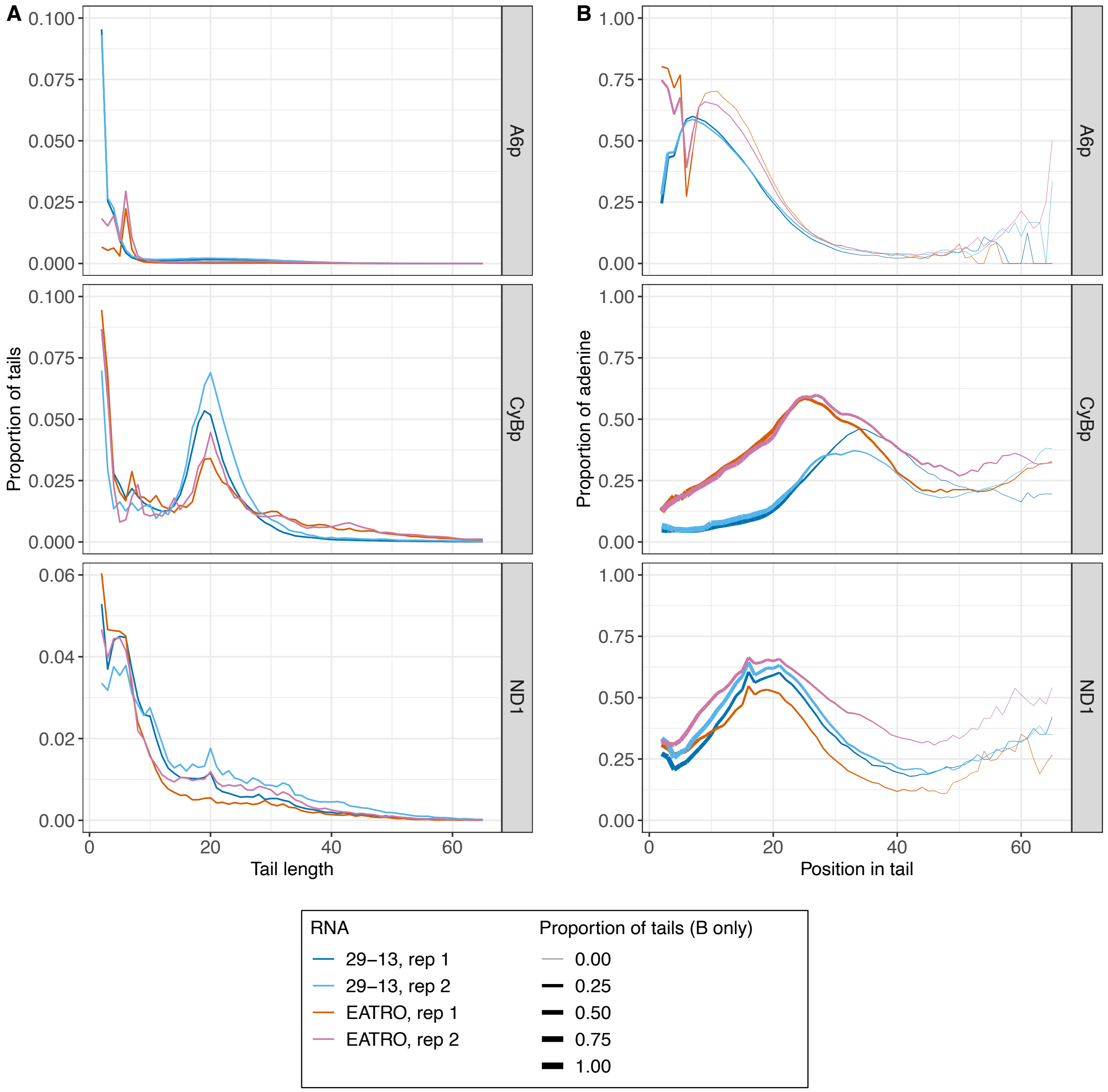


**Figure S3.** Non-encoded appended 3' nucleotide tails incorporated into *Trypanosoma brucei* mitochondrial circularized mRNA (circRNA) have similar characteristics in 29-13 and EATRO164 cell lines. Biological duplicate populations of 29-13 circRNA tails are shown in dark and light blue and tail populations from EATRO164 (EATRO) circRNA are shown in red and pink. **(A)** Tail length population density curves for three mitochondrial transcripts. **(B)** Population density curves of proportions of nucleotides in aggregate that are adenine at each tail position for each tail for three mitochondrial transcripts. The thickness of the line represents the proportion the tail population that is long enough to contribute to the data at each nucleotide position. As the lines becomes thinner, adenine content data is supported by fewer total tails because very few long tails are represented in the population. Data is shown starting from the second nucleotide in the tail to accommodate imprecision in tail cutoff determination.

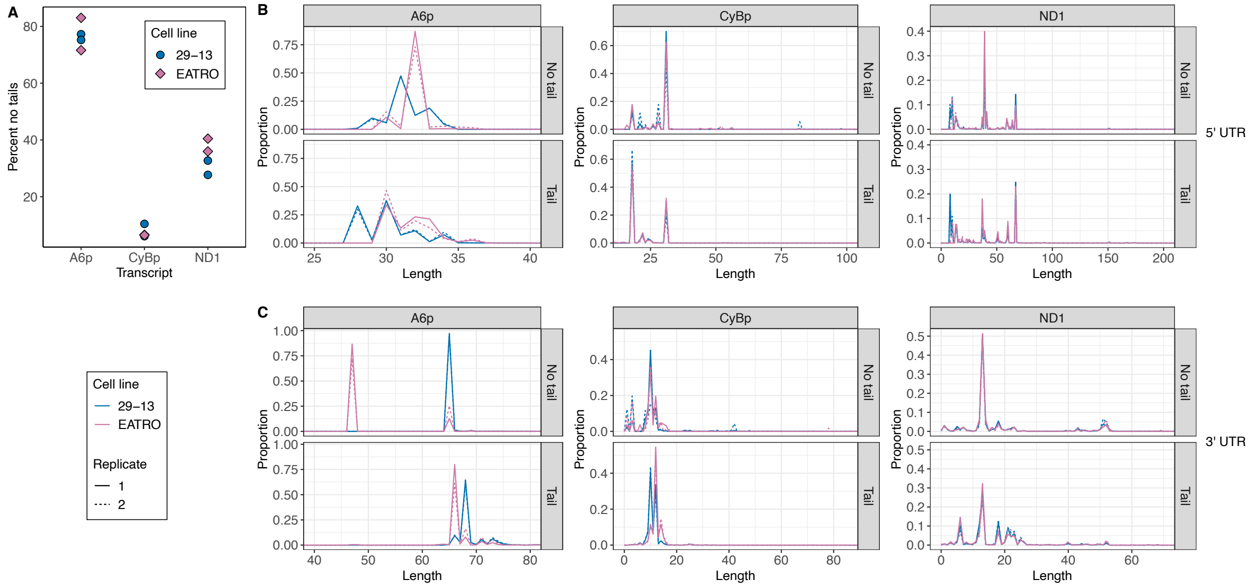


**Figure S4.** *Trypanosoma brucei* mitochondrial circularized mRNAs (circRNAs) in 29-13 and EATRO 164 cell lines have similar 5' termini and percent of molecules in which non-encoded 3' nucleotide tails were appended prior to circularization. **(A)** Percent of sequenced 3'-5' junction reads that do not possess untemplated A or U additions between the ligated 3' and 5' UTR regions. **(B)** 5' UTR lengths of circRNA in 29-13 (blue) and EATRO164 (EATRO) populations (pink) for each transcript. Populations of 3'-5' junctions that contain nontemplated tail sequence are analyzed separately from populations that do not. (C) 3' UTR lengths of circRNA 29-13 and EATRO164 populations for each transcript. 5' UTRs lengths are counted in circularized molecules starting with the first nucleotide after the tail ends until the first nucleotide of the start codon. 3' UTRs are counted so that the last nucleotide after the stop codon is position +1. Replicate 1, solid line; replicate 2, dotted line.
